# Paracetamol/acetaminophen hepatotoxicity: new markers for monitoring the elimination of the reactive N-Acetyl-p-benzoquinone imine

**DOI:** 10.1101/2023.04.28.538718

**Authors:** Eva Gorrochategui, Marc Le Vee, Habiba Selmi, Anne Gérard, Jade Chaker, Annette M Krais, Christian Lindh, Olivier Fardel, Cécile Chevrier, Pierre Le Cann, Gary W Miller, Robert Barouki, Bernard Jégou, Thomas Gicquel, David Kristensen, Arthur David

## Abstract

Paracetamol/acetaminophen (N-acetyl-p-aminophenol, APAP) overdose is one of the most important causes of drug-induced liver injury worldwide. Hepatotoxicity induced by APAP is mainly caused by the production of N-acetyl-p-benzoquinone imine (NAPQI), a highly reactive intermediate formed predominantly via the cytochrome P450 2E1. Here, we used human studies and *in vitro* models to demonstrate that NAPQI-derived thiomethyl metabolites identified using high-resolution mass spectrometry could serve to monitor NAPQI detoxification and elimination in patients (after intake at recommended dose or after intoxication), and to study inter-individual variability in NAPQI production. Using *in vitro* human models, we showed that these thiomethyl metabolites are directly linked to NAPQI detoxification since they are mainly formed after exposure to glutathione-derived conjugates via an overlooked pathway called the thiomethyl shunt. These long-term thiomethyl metabolites have great potential in future clinical studies in order to provide a more reliable history of APAP ingestion in case of acute intoxication or to study underlying causes involved in APAP-induced hepatotoxicity.

**One Sentence Summary:** Thiomethyl metabolites are new markers to monitor the elimination of the toxic N-acetyl-p-benzoquinone imine after therapeutic use or intoxication.

## INTRODUCTION

Despite being considered one the safest pharmaceuticals, paracetamol/acetaminophen (N-acetyl-p-aminophenol, APAP) overdose is one of the most important causes of drug-induced liver injury, that can eventually lead to acute liver failure (ALF) (*1, 2*). Data from the U.S. Acute Liver Failure Study Group registry of more than 700 patients with ALF across the US, implicates APAP poisoning in nearly 50% of all ALF. In Europe, APAP overdose was found to represent one-sixth of all-cause ALF (*2*). Overall, APAP overdose, even without suicidal intent, represents a large proportion of ALF leading to registration for transplantation (*1*).

APAP-induced hepatotoxicity is mainly caused by the production of N-acetyl-p-benzoquinone imine (NAPQI), a highly reactive intermediate formed predominantly via the cytochrome P450 2E1 (CYP2E1). NAPQI has been linked to hepatocellular injury due to its electrophilic nature and its covalent binding to cysteine molecules on liver proteins (*3*). At therapeutic doses, healthy livers detoxify NAPQI by phase II conjugation with glutathione (GSH) to form inert mercapturate and cysteine metabolites. When GSH stores are reduced due to APAP overdose, the detoxification capacity of the liver is exceeded and NAPQI accumulates, leading to hepatocytes necrosis (*4*).

NAPQI production estimated via the elimination of glutathione-derived metabolites (*i.e.*, mercapturate and cysteine conjugates) in urine is considered to be approximately 5-10% of the initial dose (*6–8*). The glutathione conjugates also form thiomethyl metabolites such as S-methyl-3-thioacetaminophen (S-CH_3_-APAP) (*9, 10*), but they are considered to be negligible (<1% of the initial dose) (*11*). We recently demonstrated using a pharmacometabolomics approach based on high-resolution mass spectrometry (HRMS) that significant signals corresponding to the conjugated forms of S-methyl-3-thioacetaminophen and its sulfoxide can be detected in urine from male volunteers after APAP administration. Moreover, the detected thiomethyl metabolites were of comparable diagnostic sensitivity as major APAP phase II conjugates, and exhibit delayed peak concentrations probably due to a late reabsorption after the enterohepatic circulation (*11*). Hence, we postulate that the aforementioned thiomethyl metabolites can improve monitoring of NAPQI detoxification and elimination in humans, with implications for studying risks and susceptibility to develop APAP hepatotoxicity.

Here, we studied the formation rates of these NAPQI-derived thiomethyl metabolites *in vivo* (men and women) after the use of APAP at recommended doses and after acute intoxication in order to understand their relevance as markers to monitor NAPQI elimination and determine the timing of APAP intake. Using a combination of observational human studies and *in vitro* (*i.e.*, liver models and bacterial strains), our findings support the relevance of these thiomethyl metabolites in future clinical settings aiming to provide a more reliable history of APAP ingestion, in particular to monitor the elimination of the toxic fraction after acute intoxication, or to assess inter-individual variability in the production of NAPQI.

## RESULTS

### Thiomethyl metabolites are consistently detected in men and pregnant women exposed to APAP

Thiomethyl metabolites such as S-CH_3_-APAP are produced from NAPQI-glutathione conjugates via the thiomethyl shunt, which involves the activity of a cysteine-conjugate beta lyase (CCBL) to cleave the C-S bond of cysteine conjugate, such as APAP-Cysteine (APAP-Cys) (*12*) (Fig. 1). This reaction yields ammonia, pyruvate and a reactive thiol (SH-APAP) which can be subsequently methylated with an active form of methionine (*13*). To date, these thiomethyl metabolites of APAP have been neglected as they were estimated to account for less than 1% of the initial dose in humans (*10, 14*).

**Figure 1.**
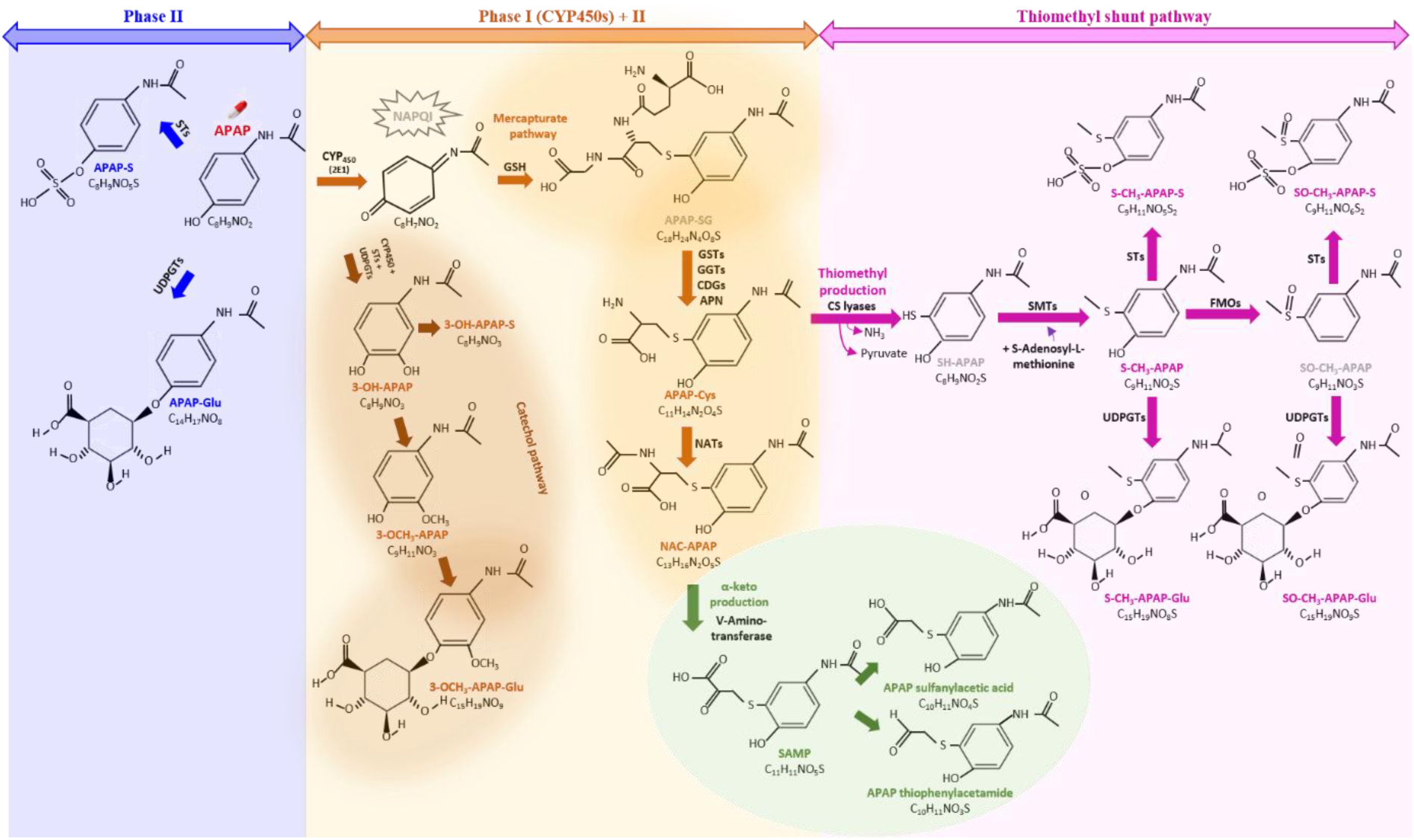
Pathways involved in human APAP metabolism (including the thiomethyl shunt recently highlighted using HRMS based non-targeted analyses (*11*)) and structures of the main APAP metabolites detected in negative mode using a UHPLC-ESI-qTOF. All metabolites were annotated based on MS2 confidence levels using a reference standard when available (see Table S9). APAP metabolites in grey are known precursors but not detected during this experiment because of their lability or low levels. APAP: acetaminophen; APAP-S: APAP sulfate; APAP-Glu: APAP glucuronide; NAPQI: N-acetyl-*p*-benzoquinone imine; 3-OH-APAP: 3-hydroxyacetaminophen; 3-OH-APAP-S: 3-hydroxyacetaminophen sulfate; 3-OCH3-APAP: 3-methoxyacetaminophen; 3-OCH3-APAP-Glu: 3-methoxyacetaminophen glucuronide; APAP-SG: acetaminophen glutathione; APAP-Cys: 3-(Cystein-S-yl)acetaminophen; NAC-APAP: Acetaminophen mercapturate; NAC-O-APAP: Acetaminophen mercapturate sulfoxide; SH-APAP: 3-mercaptoacetaminophen; SH-APAP-Glu: 3-mercaptoacetaminophen glucuronide; SH-APAP-S: 3-mercaptoacetaminophen sulfate; S-CH3-APAP: S-methyl-3-thioacetaminophen; S-CH3-APAP-S: S-methyl-3-thioacetaminophen sulfate; S-CH3-APAP-Glu: S-methyl-3-thioacetaminophen glucuronide; SO-CH3-APAP: S-methyl-3-thioacetaminophen sulphoxide; SO-CH3-APAP-S: S-methyl-3-thioacetaminophen sulphoxide sulphate; SO-CH3-APAP-Glu:: S-methyl-3-thioacetaminophen sulphoxide glucuronide; STs: sulfotransferases; UDPGTs: uridine diphosphoglucuronyltransferases; GSTs: glutathione *S*-transferases; GGTs: gamma glutamyltransferases, CGDs: cysteinylglycine dipeptidases; NATs: N-acetyltransferases; CS lyases: Cysteine S-conjugate β-lyases; SMTs: S-methyl-transferases; FMOs: flavin-dependent monooxygenases; APN: aminopeptidase N.

We recently demonstrated using HRMS-based analytical methods that significant signals can be observed for the conjugated forms of S-CH_3_-APAP (*i.e.*, S-methyl-3-thioacetaminophen sulfate [S-CH_3_-APAP-S]) and its sulphoxide (S-methyl-3-thioacetaminophen sulphoxide [SO-CH_3_-APAP-S]) in urine from male volunteers who ingested 1 g of APAP (*11*). Delayed formation and excretion rates were observed for these thiomethyl metabolites (peak time at 24h after APAP intake) compared to conventional APAP metabolites (peak time 2-4h after intake) (Fig. 2A) (*11*).

**Figure 2.**
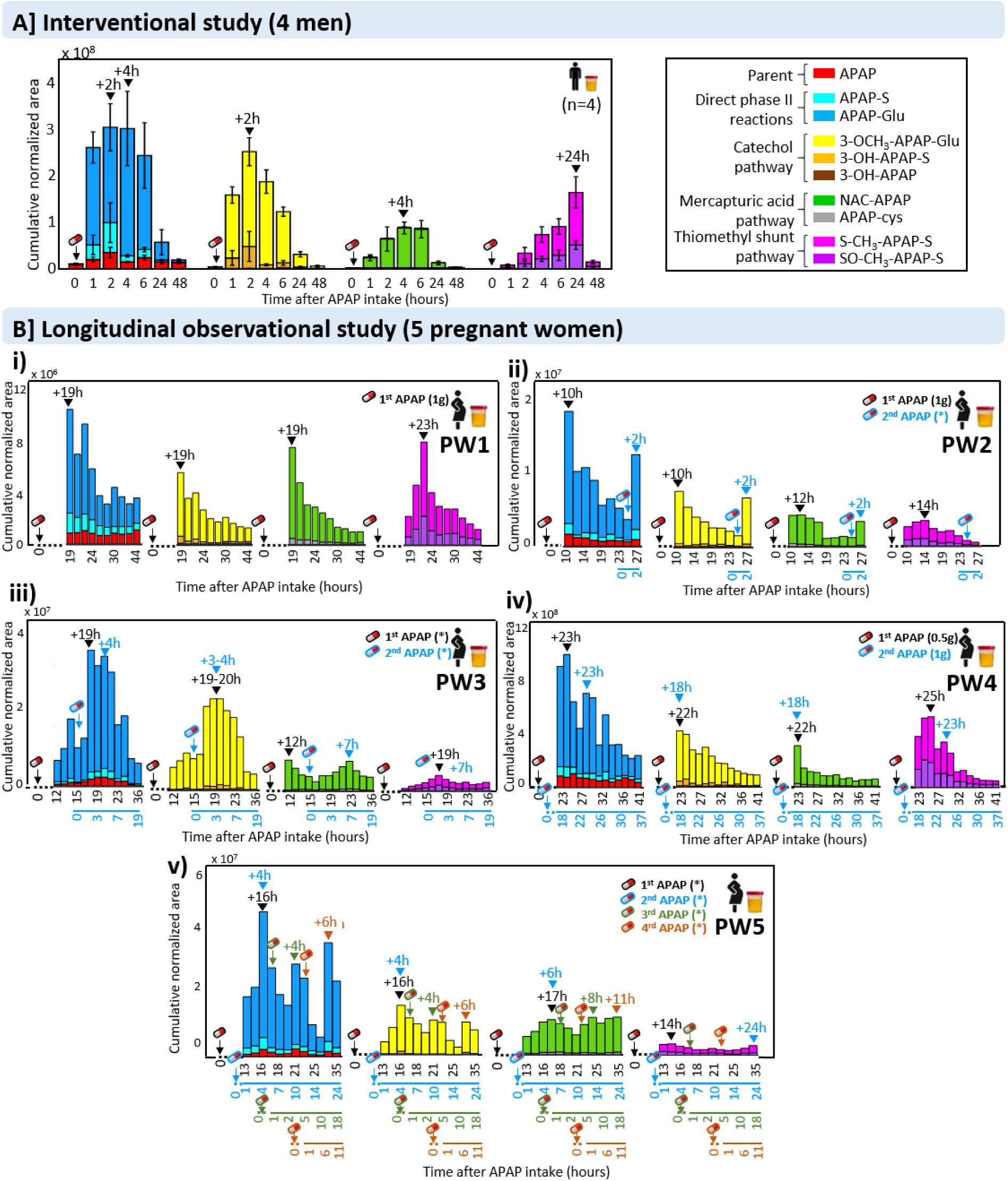
Kinetics of formation of free APAP, phase II metabolites (APAP-Glu and APAP-S), catechol metabolites (3-OCH3-APAP-Glu, OH-APAP-S, and OH-APAP), glutathione derived metabolites (NAC-APAP and APAP-Cys) and thiomethyl metabolites (S-CH3-APAP-S and SO-CH3-APAP-S) detected using UHPLC-ESI-QTOF-MS analyses in A) urine of 4-men (mean ± SEM) before intake (baseline, n=12) and at different time points after intake of 1g APAP (interventional study, (*11*)) and B) urine of 5 pregnant women (PW) from the longitudinal observational study (plots i to v) at different time points after intake (0h) of ≥1 APAP doses (represented with different colors in the figure). In all the plots, the integrated peak areas of individual markers were normalised on the total peak area (TIC). Medications containing APAP and other co-administration were taken by the 5 pregnant women: PW1: containing APAP (1x doliprane 1 g), co-administration (1x betamethasone, 2x Vicks inhaler); PW2: containing APAP (2x paracetamol), co-administration (3x esomeprazole, 2x plaquenil 400 mg and 2x leucocetirizine 5 mg); PW3: containing APAP (2x doliprane), co-administration (2x omeoprazol, 3x spasfon, 1.5x dolormyl); PW4: containing APAP (1x doliprane 0.5 mg, 1x doliprane 1g), co-administration (3x mupirocine 2% pomade, 1x mupirocine); PW5: containing APAP (6x dafalgan), co-administration (2x levothyrox, 4x timoferol, 2x magnesium, 2x gestarelle, 2x esomeprazole) (for more information see Table S3). Annotation: To facilitate reading, for PW5, despite the intake of 6 APAP doses, only the time periods of the first 4 doses of APAP have been represented in the figure.*= unknown amount of APAP.

In the present study, we monitored and compared the formation of S-CH_3_-APAP-S and SO-CH_3_-APAP-S in urine samples collected from five pregnant women (PW) in their 2^nd^ trimester of pregnancy, up to 2 days after a single or repeated APAP intake (n=57). Pregnancy has previously been shown to be associated with increased oxidative metabolism (*15*). Furthermore, APAP is the most commonly used pharmaceutical during pregnancy (worldwide, more than 50% of pregnant women are estimated to use APAP (*16, 17*)), and the main cause of drug overdose during this period (*18*). Areas under the curve (AUCs) based on normalised areas obtained from LC-HRMS were calculated for all APAP metabolites (table S1) previously annotated using MS/MS structural elucidation (*11*) (see Fig.1) to assess differences in the diagnostic signals produced by this analytical technique.

We confirmed that significant signals for S-CH_3_-APAP-S and SO-CH_3_-APAP-S could be detected in all PWs (AUC for combined metabolites represented an average 13±9% of all AUCs, table S1, Fig. 2B), and that inter-individual variabilities were evident (combined AUCs for S-CH_3_-APAP-S and SO-CH_3_-APAP-S ranged from 5-6% (PW 5 and 3) to 21-24% (PW 4 and 1) of all AUCs (table S1)). Overall, the formation of the phase II glucuronide conjugates was the dominant pathway in all PWs in terms of signals observed with our HRMS-based method (average AUCs of 48 ± 5%), followed by those related to the catechol metabolites (average AUCs of 20 ± 6%), the mercapturic pathway-related metabolites (average AUCs of 15 ± 5%), and then the thiomethyl conjugates. Altogether, the AUCs based on signals observed for NAPQI-derived metabolites (*i.e*., from the mercapturic pathway and the thiomethyl shunt combined) represent up to 28% of all AUCs. The signals from thiomethyl metabolites can even be dominant in some cases compared to those observed from the mercapturic pathways (*e.g.*, for PW1 and PW4), showing the importance of these thiomethyl metabolites to monitor more accurately the total excretion of the toxic NAPQI using current LC-ESI-MS-based methods.

We also confirmed the potential of S-CH_3_-APAP-S and SO-CH_3_-APAP-S to identify inter-individual variability in hepatotoxic NAPQI formation, in particular when they are combined with mercapturic metabolites (their combined AUCs varied by twice as much, from 19 to 41% of all AUCs for the 5 PWs). The inter-individual variabilities observed for the formation of S-CH_3_-APAP-S and SO-CH_3_-APAP-S in PWs are similar to the ones observed in the interventional study in men (Fig. 2A, table S2) (*11*). The inter-individual variabilities observed for the 5 PW could be related to the individual intrinsic ability to generate NAPQI-derived metabolites (genetic factors), and/or could be influenced by drug co-administration (table S3) or other environmental factors (including other xenobiotics (*19*)) involved in CYP2E1 induction (*e.g.*, nutrition and fasting) (*20*).

A delayed concentration peak was observed for thiomethyl metabolites compared to others in the PWs (from 2 to 20h, Fig. 2B), even though the delay was more variable and sometimes lower than those observed in the interventional study (20-22h, Fig. 2A). The lack of control over the initial intake and other factors like co-administration or repeated intakes may have influenced the kinetics observed in this observational study with PWs. This delayed appearance is consistent with the fact that the formation of S-CH_3_-APAP-S and SO-CH_3_-APAP-S via the thiomethyl shunt is thought to be limited by prior chemical and biological steps (*i.e.*, the formation of glutathione-derived metabolites and their reabsorption after the enterohepatic circulation as observed in rodents) (*11, 21*), as opposed to the direct excretion in urine for other metabolites.

Significant AUCs based on diagnostic signals similar to those observed for the glucuronide conjugates were observed for catechol metabolites, and in particular for 3-methoxyacetaminophen glucuronide (3-OCH_3_-APAP-Glu) (AUC ranged from 18% to 33% of all AUCs) (table S1). Comparisons of AUCs for oxidative catechol and NAPQI-derived metabolites do not suggest a difference in the production of oxidative metabolites between men and pregnant women in their 2^nd^ trimester.

### Thiomethyl metabolites inform on the timing of intakes and identify a hyper NAPQI-producer

We then monitored the presence of these thiomethyl metabolites in a cross-sectional study using urine collected from 10 PWs who reported APAP use and 10 PW who did not (based on questionnaires). In addition to questionnaires and urinary LC-HRMS profiling, we also used conventional targeted LC-MS/MS to identify accurately PWs with urinary APAP concentrations within therapeutic doses.

The 10 PWs that did not report APAP consumption had basal APAP urinary concentrations (median of 125 ng/ml, table S4) (Fig. 3A) similar to urinary environmental concentrations already observed in the general population (medians of 61.7 ng/ml and 100 ng/ml in the German and Danish populations, respectively) (*11, 22, 23*). The sources of environmental exposure to APAP have been linked to APAP precursors such as aniline and 4-aminophenol (two large-volume intermediates in industrial processes), and/or indirect APAP exposure through environmental sources (*17, 22, 24*). In these PWs, thiomethyl metabolites could only be detected in one PW (*i.e.*, PW3). The presence of low detectable levels of S-CH_3_-APAP-S and SO-CH_3_-APAP-S in PW3 could be related to a higher APAP environmental exposure (*i.e.* 1.4·10^3^ ng/ml as opposed to the median of 125 ng/ml), or more simply to a recall bias.

**Figure 3.**
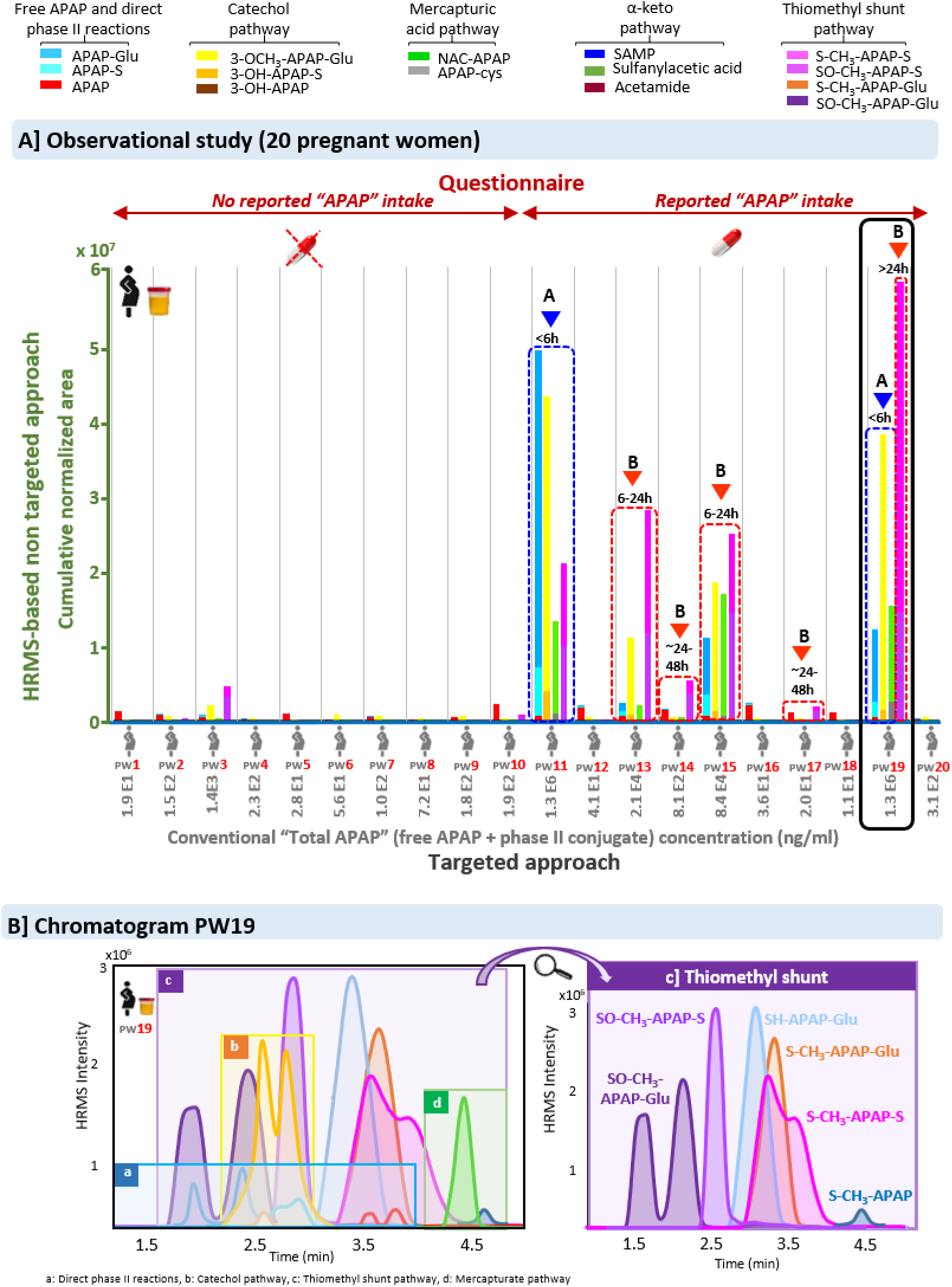
A] HRMS-based profiles of APAP metabolites in urine detected in the cross-sectional study performed with 20 pregnant women, including 10 women who declared “no APAP intake” versus 10 women who declared “APAP intake”. “Total” APAP concentrations (free APAP + phase II conjugate) after enzymatic deconjugation using LC-MS/MS were measured for each pregnant woman and are represented in ng/ml. A corresponds to HRMS-based profiles where Phase II + catechol > Thiomethyl shunt, B corresponds to HRMS-based profiles where Thiomethyl shunt > Phase II + catechol. B] UHPLC chromatogram of APAP metabolites in pregnant woman 19 that shows high levels of thiomethyl metabolites.

For PWs who reported APAP consumption, the comparison of results from targeted and non-targeted LC-HRMS-based analyses demonstrate the ability of thiomethyl metabolites to inform on the timing of APAP ingestion (*i.e.*, based on the fact that thiomethyl metabolites have delayed kinetics compared to other conventionally investigated APAP metabolites). Hence, urinary targeted analyses and HRMS-based profiles allowed to distinguish PWs who had urine collected <6h after APAP ingestion (*e.g.*, PW11) (Fig. 3A). This PW had APAP urinary concentrations within those generated by therapeutic ranges (1.3·10^6^ ng/ml) and an HRMS-based profile matching what was observed during the first 6h of the 4-men interventional study (*i.e.*, direct phase II conjugates and catechols’ signals higher than those of thiomethyl metabolites) (*11*). Likewise, urinary targeted analyses and HRMS-based profiles allowed to distinguish PWs for whom urine collection was performed between 6 and 24h after APAP ingestion (*e.g.*, PWs 13 and 15). These PWs had urinary APAP concentrations below therapeutic ranges but higher than basal concentrations (2.1·10^4^ – 8.4·10^4^ ng/ml, respectively) while HRMS-based metabolite profiles (*i.e.*, thiomethyl metabolites signals higher than direct phase II conjugates and catechols) are consistent with the profiles observed 24h after APAP intake in the interventional study.

Furthermore, LC-HRMS-based analyses also provided additional observations compared to targeted analyses and/or the questionnaires. In particular, the presence of thiomethyl metabolites helped detecting intentional APAP intake that occurred ∼24-48 h before sampling. At this time point, environmental basal APAP urinary concentrations (*11, 22*) are interfering with the assessment of APAP therapeutic use since APAP and its glucuronide conjugate have nearly disappeared from urine. However, the presence of S-CH_3_-APAP-S and SO-CH_3_-APAP-S, still detectable using LC-HRMS at sufficient levels, allows to confirm a previous intake >24h before urine collection (*e.g*. PWs 14 and 17).

We also observed one peculiar LC-HRMS-based metabolite profile for PW19 who reported APAP use, which could serve to identify individuals that had repeated APAP intakes. PW 19 had APAP urinary concentrations within the range of those related to therapeutic use (1.3·10^6^ ng/ml) while both levels of short-term metabolites (*e.g.*, 3-OCH_3_-APAP-Glu) and long-term metabolites (*i.e.*, S-CH_3_-APAP-S and SO-CH_3_-APAP-S) detected with LC-HRMS were high. This combination would suggest that urine was collected after a course of both recent (<6h) and late intake (>24h). Furthermore, unusually high levels of S-CH_3_-APAP-S and SO-CH_3_-APAP-S but also other conjugates and intermediates forms of thiomethyl metabolites were found. This is the case of conjugated glucuronide forms (*i.e.*, S-methyl-3-thioacetaminophen glucuronide [S-CH3-APAP-Glu], S-methyl-3-thioacetaminophen sulphoxide glucuronide [SO-CH3-APAP-Glu], the intermediate conjugated 3-thioacetaminophen glucuronide [SH-APAP-Glu], and non-conjugated S-CH_3_-APAP, Fig. 3B). The high levels of metabolites related to the thiomethyl shunt suggest an excessive production of NAQPI-derived glutathione conjugates (unseen so far in the other individuals). However, it is unclear if this abundance of thiomethyl derivates is related to a specific susceptibility of this individual to generate NAPQI metabolites, to the pregnancy itself (as sex steroids could be involved in the regulation of CYP2E1 (*25*)) or to an accumulation of these metabolites in urine over time due to repeated intakes.

### Thiomethyl metabolites allow monitoring the elimination of the toxic fraction after acute intoxication

We then monitored the presence of S-CH_3_-APAP-S and SO-CH_3_-APAP-S in blood samples collected from individuals who were admitted to hospital for suspicion of acute APAP intoxication (n=13, 11 women and 2 men). Based on hospital protocol, the antidote (N-acetylcysteine, NAC, weight-based dosage) was administered in 8 out of 13 patients, sometimes at several time points. The decision to administer the antidote was based on measurements of blood serum APAP concentrations performed using absorption spectrophotometric measurement COBAS® assays (Roche Diagnostic®) and clinical expertise of emergency doctors. All individuals we selected had repeated collections of blood samples (n=12 with 2 repeated blood samples and n=1 with 3 repeats) in order to study the formation of S-CH_3_-APAP-S and SO-CH_3_-APAP-S signals over time as an indicator of NAPQI elimination.

Blood serum APAP concentrations measured in these patients ranged from 4 to 284 mg/L (median 52 mg/L), and correlated positively with APAP signals measured with the HRMS-based method (r^2^= 0.89, table S7). The repeated blood samples collected allowed observing significant reductions in APAP concentrations over time (*i.e.*, from 24% of initial concentration up to 97% for the repeated sample, see table S7) for 11 out of 13 patients.

Our results confirmed that the reduction of APAP concentration in blood observed over time in intoxicated patients was always associated with an increase in thiomethyl metabolites signals (mainly as S-CH_3_-APAP-S as observed in the intervention study) (Fig. 4B). In 4 out of 11 patients (*i.e.*, I, J, K and M), the signals observed for thiomethyl metabolites were equivalent or higher than the ones observed for free APAP, indicating that a high proportion of reactive NAPQI was generated in these patients, and then reabsorbed into the systemic circulation before being eliminated. For these 4 patients with equivalent or higher thiomethyl metabolite signals than the ones observed for APAP, the timing of the second blood collection was always >11h (between 11h and 42h), and the percentage of APAP reduction >90% (see table S7). Signals of thiomethyl metabolites were always low for blood samples collected less than 6h apart, showing that elimination of NAPQI (either after conjugation to glutathione or the antidote) is also subject to a delay in case of acute intoxication.

**Figure 4.**
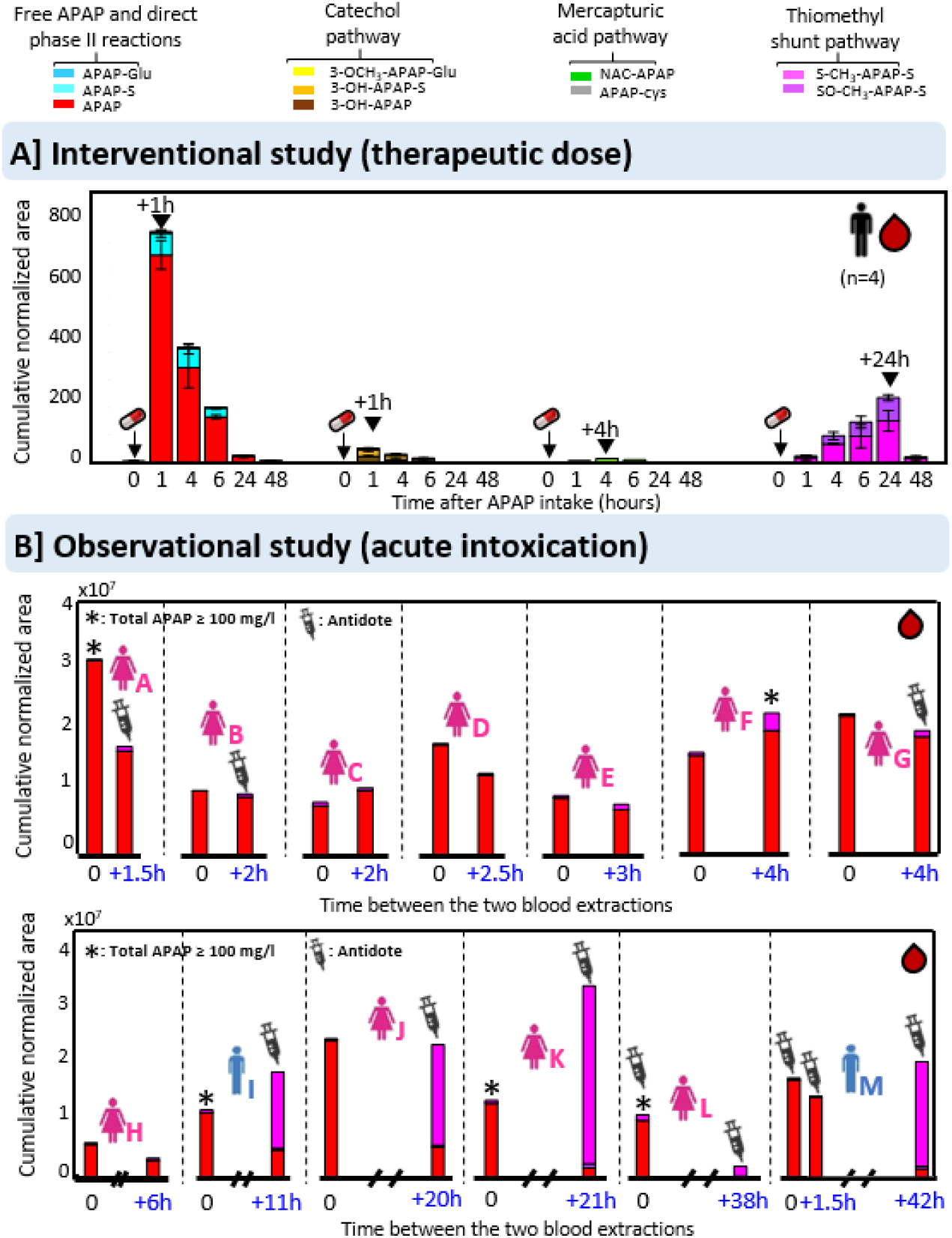
A) Kinetics of formation of free APAP, phase II metabolites (APAP-Glu and APAP-S), catechol metabolites (3-OCH3-APAP-Glu, OH-APAP-S, and OH-APAP), glutathione derived metabolites (NAC-APAP and APAP-Cys) and thiomethyl metabolites (S-CH3-APAP-S and SO-CH3-APAP-S) detected using non-targeted screening based on UHPLC-ESI-QTOF-MS analyses in blood plasma of 4-men (mean ± SEM) before intake (baseline, n=12) and at different time points after intake of 1g APAP (interventional study, (*11*)) and B) Distribution profiles of free APAP and thiomethyl shunt metabolites in the observational study performed with serum samples of patients under acute APAP intoxication after consecutive sample intakes. Patients which were treated with N-acetylcysteine (NAC) antidote are indicated in the figure.

Hence, we confirmed that both thiomethyl metabolites also provide significant signals in blood in addition to urine (Fig. 4A), indicating the potential of these conjugates to serve as relevant clinical markers when blood samples are available, as it is in the case of acute intoxication, to monitor the elimination of the toxic NAPQI.

### The liver produces high levels of thiomethyl metabolites from glutathione conjugates

We then used *in vitro* liver models (HepaRG-HRG-and primary human hepatocytes-PHH) to provide a better understanding of the formation of these thiomethyl metabolites (in particular to identify their precursors), and a better understanding of the site where the biotransformation occurs. The biotransformation that leads to the delayed formation of thiomethyl metabolites via the thiomethyl shunt pathway has been shown to occur in rodents via the enterohepatic circulation and biliary excretion of the glutathione-derived conjugates into the intestine (*9*). However, it remains unclear whether the thiomethyl shunt involved in the formation of S-CH_3_-APAP-S and SO-CH_3_-APAP-S is preferentially occurring in the liver (before excretion of thiomethyl metabolites in the bile), or via the microbiota (*26, 27*).

We exposed HRG cells (4 independent experiments) and PHH (3 independent experiments, from 3 independent donors) to APAP and APAP-cysteine (APAP-Cys), separately, at 50 µg/ml. APAP was used to determine whether the cells could produce thiomethyl metabolites *in vitro* from APAP, which implies multiple successive steps, and the involvement of several enzymes (*i.e.*, CYP2E1 > glutathione S-transferases > 3 γ-glutamyltransferase > dipeptidases > CCBL > thiomethyltransferase). The second experiment involved exposure to APAP-Cys as it is known as the main precursor of CCBL in the thiomethyl shunt pathway (*13*). A 24h exposure time was selected for both HRG cells and PHH as the optimal time of exposure, as indicated by kinetic studies using several time points (Fig. 5).

**Figure 5.**
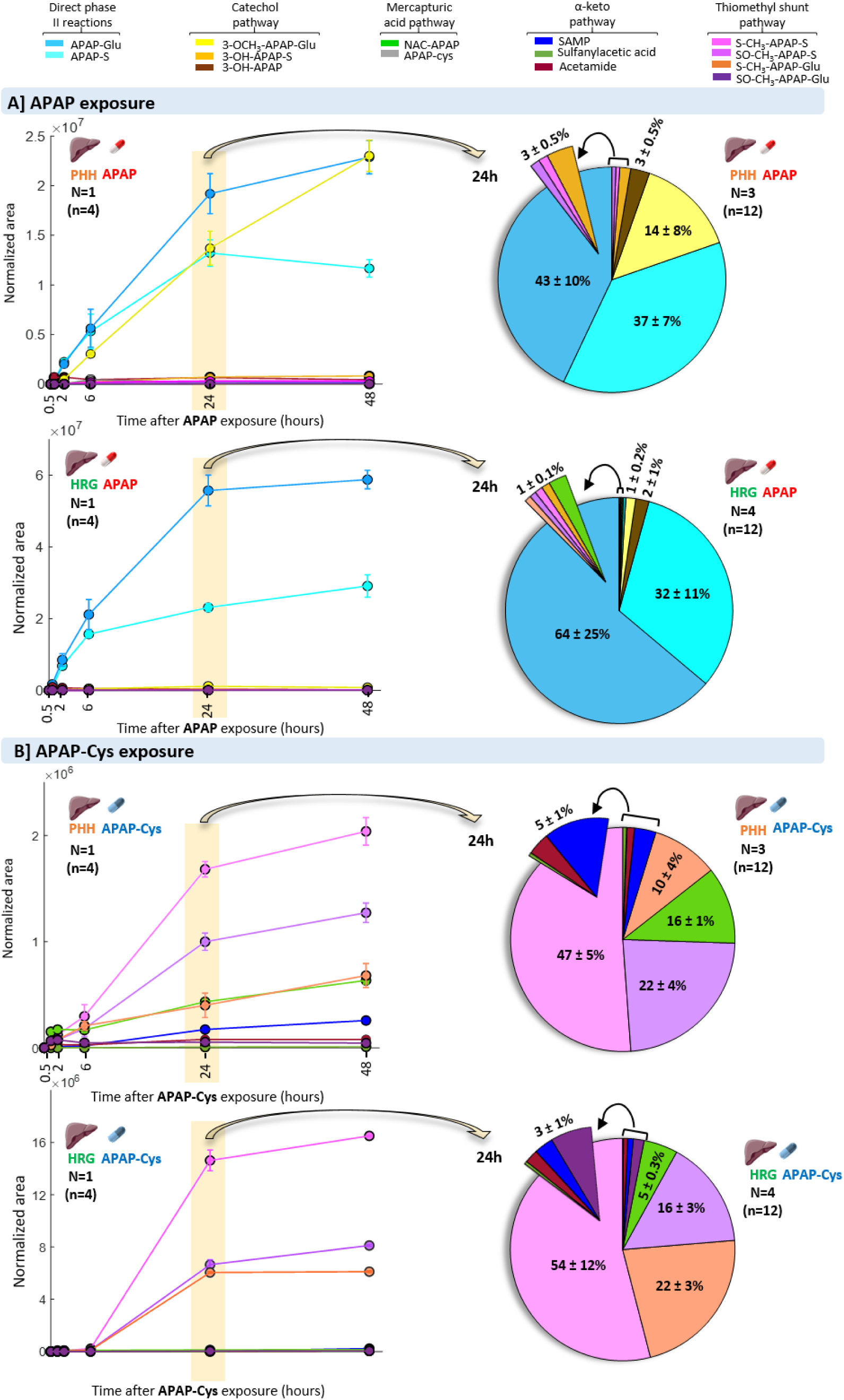
Left side: Kinetics of formation of APAP metabolites primary human hepatocytes (PHH) and HepaRG cells (HRG) exposed to A] free APAP and B] to APAP-Cys at a concentration of [50 µg/ml] at different time points: 0, 0.5, 2, 6, 24 and 48h. Right side: Pie charts to illustrate the metabolite proportions (mean ± SEM) at 24h of exposure. N=number of independent experiments, n=total number of technical replicates within a given experiment.

All metabolites detected *in vivo* could be detected after exposure to APAP for 24h in HRG cells and PHH. In both models, levels of thiomethyl metabolites were low (normalised area of S-CH3-APAP-S and SO-CH3-APAP-S represent less than 3 and 1% in PHH and HRG of all detected metabolite’s areas, respectively) compared to direct phase II conjugates (Fig. 5A). They, therefore, differed from the higher thiomethyl signal observed *in vivo* (see Fig 2 and 3). As mentioned previously, the formation of thiomethyl metabolites is dependent on NAPQI production by CYP2E1. Hence, the low levels of thiomethyl metabolites observed after APAP exposure are probably associated to the limited cytochrome P450 capabilities of *in vitro* models to generate NAPQI metabolites via CYP2E compared to *in vivo* hepatocytes. This seems to be consistent with the fact that higher levels of thiomethyl metabolites were observed in PHH than in HRG cells since CYP2E1 activity in primary human hepatocytes is known to be higher than in HRG cells (*28*). In agreement with this, higher levels of the catechol oxidative metabolite 3-OCH_3_-APAP-Glu were also observed in PHH compared to HRG cells.

We then exposed HRG cells and PHH to APAP-cysteine for 24h to study their ability to generate thiomethyl metabolites. We confirmed here that APAP-cysteine is an important and direct precursor of thiomethyl metabolites since the conjugated forms S-CH_3_-APAP and SO-CH_3_-APAP were the dominant metabolites detected in both models. High levels of both sulfate conjugates (as observed *in vivo*) but also glucuronide conjugates (*i.e.*, S-methyl-3-thioacetaminophen glucuronide [S-CH_3_-APAP-Glu]) could be detected (normalised total area of S-CH_3_-APAP-S and SO-CH_3_-APAP-S represent 69 and 76% of all metabolites in PHH and HRG, respectively). Hence, we showed that both liver models produce CCBL enzymes that have the ability to cleave the C-S bond of cysteine conjugates, which tends to demonstrate the involvement of the liver in the formation of these thiomethyl metabolites.

Furthermore, HRMS-based non-targeted analyses using MS/MS structural elucidation allowed to identify other relatively minor metabolites related to APAP-Cys such as S-(5-acetylamino-2-hydroxyphenyl)mercaptopyruvic acid (SAMP), an APAP metabolite potentially formed by the transamination reaction of APAP-cysteine (see Fig.1), previously observed in mouse (*29*). The conversion of a cysteine S-conjugate to the corresponding alpha-keto acid occurring by transamination with a suitable alpha-keto acid acceptor has already been observed for other compounds conjugated to NAC (*27*). Two other minor metabolites were annotated as N-(4-hydroxy-3-((2-oxoethyl)thio)phenyl)acetamide and 2-(5-Acetamido-2-hydroxyphenyl)sulfanylacetic acid. These minor metabolites could also be detected after APAP exposure (but at lower levels), and *in vivo* in the men and PWs, but at much lower levels than the thiomethyl metabolites (AUCs for SAMP in the men and PW correspond to less than 4% of all combined AUCs *in vivo,* see tables S1 and S2). Furthermore, the peak concentrations of these minor metabolites were observed earlier than those of thiomethyl metabolites (between 2 and 6 h as opposed to 24h) (Fig. S1), suggesting that they are not involved in the thiomethyl shunt.

We then compared the production of thiomethyl metabolites after exposure to APAP, and 3 of its potential direct precursors: the glutathione conjugate (APAP-SG), APAP-Cys, and NAC-APAP for 24h in HRG cells and PHH (Fig. 6). All glutathione-derived metabolites produced significantly higher levels of S-CH_3_-APAP-S and SO-CH_3_-APAP-S than APAP (p<0.001 in HRG cells and PHH, respectively), which is in agreement with our previous observations. As for the glutathione-derived metabolites, APAP-Cys and APAP-SG produced higher and similar levels of S-CH_3_-APAP-S and SO-CH_3_-APAP-S compared to NAC-APAP. The thiomethyl shunt involves the activity of CCBL to cleave cysteine S-conjugates, and subsequent methylation with an active form of methionine to produce thiomethyl metabolites (*9, 21*), meaning that APAP-SG and NAC-APAP were probably biotransformed to APAP-cysteine beforehand. Interestingly, NAC-APAP produced high levels of the SAMP and N-(4-hydroxy-3-((2-oxoethyl)thio)phenyl)acetamide compared to APAP-SG and APAP-cysteine. Such difference was more evident in HRG cells, where the production of the latter metabolites by APAP-SG and APAP-Cys was almost non-existent, but less significant in PHH, where there was a significant production of SAMP by APAP-SG (ANOVA’s *p<0.001*).

**Figure 6.**
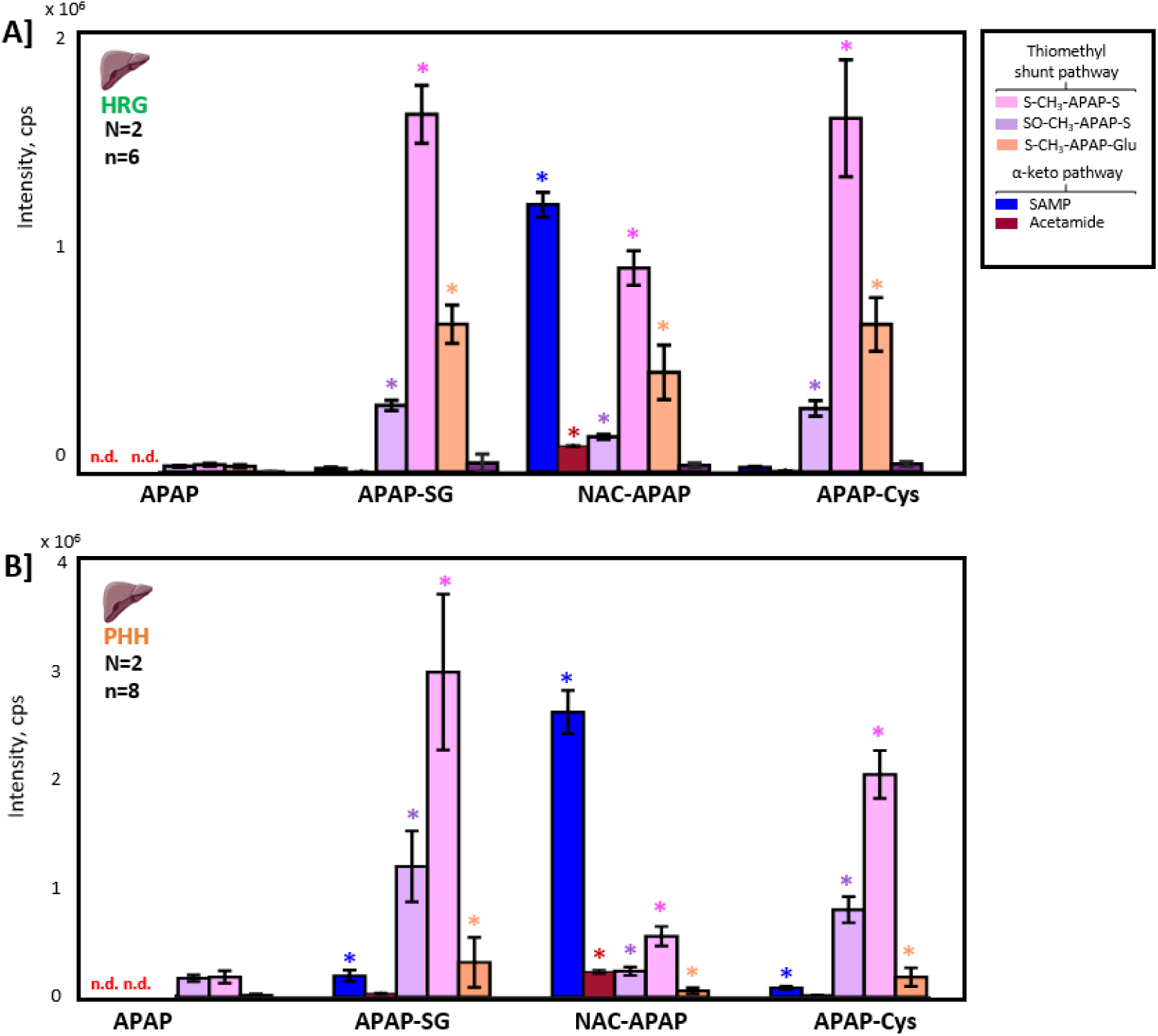
Concentrations of the thiomethyl metabolites (mean ± SEM) related to the thiomethyl shunt produced by A] HepaRG cells (HRG) and B] primary human hepatocytes (PHH) after exposure to APAP, APAP-SG, NAC-APAP and APAP-Cys at 50 µg/ml. One-way ANOVA followed by multiple comparisons test is used to indicate statistical differences between APAP-SG, NAC-APAP, and APAP-Cys treatments versus APAP (\**p<0.001*). N=number of independent experiments, n=total number of replicates.

These results confirm the ability of the liver to generate high levels of thiomethyl metabolites from all glutathione-derived metabolites. These results also challenge the fact that APAP-cysteine and NAC-APAP are the main end products of NAPQI elimination, and demonstrate the important role of the thiomethyl shunt in the detoxification of this reactive metabolite.

### Selected enteric bacteria formed very low levels of thiomethyl metabolites

The contribution of the gastrointestinal microbiota to produce thiomethyl metabolites, after biliary excretion has been previously studied in germ-free and neomycin-treated mice. The results suggested that the gut microbiota could play an important role in the C-S cleavage of APAP-Cys to form S-CH_3_-APAP-S and SO-CH_3_-APAP-S (*30*). In agreement with that, several studies have shown that many enteric bacteria contain CCBL (*26, 27*).

Here, we selected five bacterial strains representative of different phyla colonizing the human gut (*i.e., Agathobacter rectalis, Phocaeicola vulgatus, Bifidobacterium longum, Escherichia coli,* and *Enterococcus faecalis*), and that previously displayed CCBL activity for some of them (*26, 27*). The bacterial strains were exposed to APAP and APAP-Cys, separately, at 50 µg/ml for 24h in anaerobic conditions.

In these conditions, the exposure of the bacterial strains to APAP generated only very low levels of direct phase II conjugates (APAP-Glu and APAP-S) and the catechol oxidative metabolite 3-OCH_3_-APAP-Glu compared to the levels that were observed in liver models (Fig. 7). As opposed to the liver models, exposing the bacterial strains to APAP-Cys, the main precursor of CCBL, did not generate the formation of thiomethyl metabolites S-CH_3_-APAP-S and SO-CH_3_-APAP-S. However, the SAMP, N-(4-hydroxy-3-((2-oxoethyl)thio)phenyl)acetamide and 2-(5-Acetamido-2-hydroxyphenyl)sulfanylacetic acid metabolites, observed in the liver models and *in vivo,* could be generated in presence of all bacterial strains, and in some cases at higher levels compared to the liver models (*i.e.,* higher production of SAMP by *E.coli* and of N-(4-hydroxy-3-((2-oxoethyl)thio)phenyl)acetamide by *E. coli* and *P. vulgatus*), suggesting a potential involvement of the microbiota in the production of this alpha-keto acid from APAP-Cys (Fig. 7).

**Figure 7.**
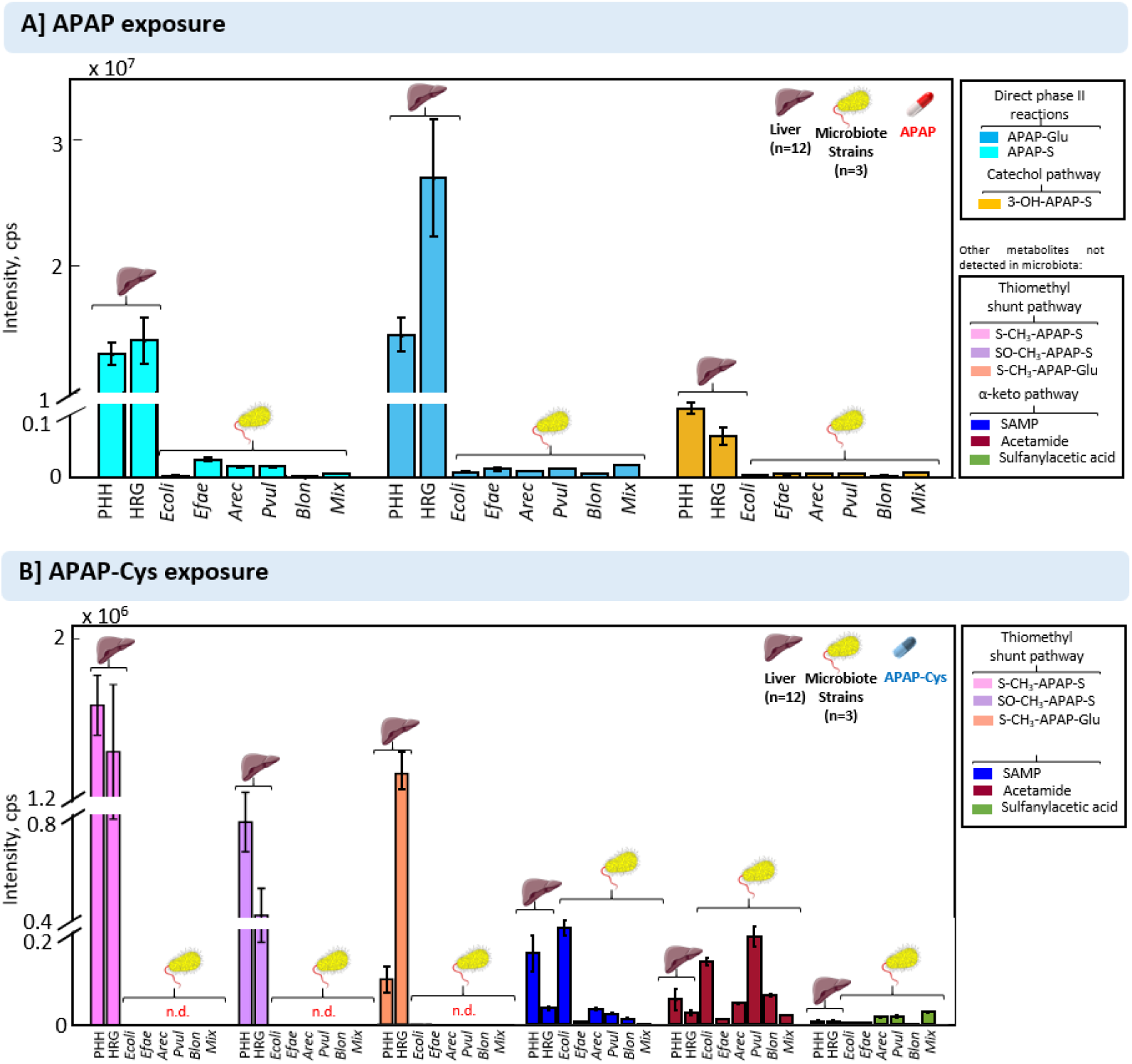
APAP metabolites produced HepaRG cells (HRG), primary human hepatocytes (PHH), and 5 types of intestinal microbiome strains after 24h exposure to A] free acetaminophen (APAP) and to B] APAP-Cys both at 50 µg/ml. Abbreviations: *Escherichia coli* (Ecoli), *Enterococcus faecalis* (Efae), *Agathobacter rectalis* (Arec), *Phocaeicola vulgatus* (Pvul), *Bifidobacterium longum* (Blon) and a mixture of the five strains (Mix). n=total number of replicates.

Since the results we observed could be related to the inability of these tests to reproduce the same conditions as the ones observed in the gastrointestinal tract for the exposure experiments (*i.e.*, in terms of T°C, pH, or even a community-based response), we then exposed APAP and APAP-Cys to lysates from all these bacterial strains obtained after cell homogenizations using sonication (*31*). Interestingly, low levels of S-CH_3_-APAP-S (but no SO-CH_3_-APAP-S) were also observed after exposure to both APAP and APAP-cysteine, suggesting that an active form of CCBL is present in these bacterial strains. No significant differences in the production of S-CH_3_-APAP-S were observed between the bacterial strains (Fig S3).

Even though these results suggest that the tested bacterial strains do not display a predominant role in the thiomethyl shunt as opposed to the liver, further experiments with the gut microbiota will be needed to confirm these results.

## DISCUSSION

APAP is an active ingredient in more than 600 prescription and non-prescription pharmaceuticals and is one of the most commonly used pharmaceuticals. The medical benefits of APAP are widely recognized. Nevertheless, APAP overdose is also an important cause of ALF worldwide. In this study, we provide evidence of the utility of new markers related to the detoxification of the toxic NAPQI that could serve to improved patient monitoring in case of APAP intoxication or to study inter-individual variability in NAPQI production. More specifically, we demonstrated that the thiomethyl metabolites S-CH_3_-APAP-S and SO-CH_3_-APAP-S are consistently detected in urine from men and pregnant women after APAP intake at therapeutic doses. We also demonstrated that high signals for S-CH_3_-APAP-S and SO-CH_3_-APAP-S can be increasingly detected in blood collected at several time points after APAP acute intoxication. Using *in vitro* liver models, we showed that these thiomethyl metabolites are directly linked to the elimination of the hepatotoxic NAPQI metabolites since S-CH_3_-APAP-S and SO-CH_3_-APAP-S are formed at high levels after exposure to NAQPI-derived metabolites (glutathione, cysteine, and mercapturate conjugates) in both primary human hepatocytes and HRG cells. We also confirmed the delayed formation of the thiomethyl metabolites compared to conventional APAP metabolites, likely related to delayed reabsorption into the systemic circulation after the enterohepatic circulation. Exposure on liver models and enteric bacteria suggests that the liver is the main site of biotransformation of these metabolites.

Based on our observations, we believe that the measurement of these thiomethyl metabolites in clinical studies could serve to provide a more reliable history of APAP ingestion in patients by studying the ratio of the short-term metabolites (*i.e.*, phase II glucuronide conjugates and conjugated methoxy APAP) over the long-term ones (*i.e.*, thiomethyl metabolites). Potential clinical applications for these markers include i) a better typing of patients after APAP acute intoxication (*i.e.*, to monitor more precisely the kinetic detoxification process through the appearance of thiomethyl metabolites); ii) a better typing of chronic users to study risks of developing hepatotoxicity at the highest recommended doses (even though this is still controversial there is accumulating evidence that the maximum recommended dose can induce mild-to-moderate hepatic cytolysis); and iii) identification of inter-individual-variability in NAPQI production and consequently identify individuals or populations with underlying predispositions (*e.g.* pre-existent induction of CYP2E1) to develop APAP-induced liver injuries. Regarding the latter application, some clinical investigations reported that non-alcoholic fatty liver disease (NAFLD), the most common chronic liver disease worldwide (*32*), could favor APAP-induced AFL after an overdose (*33, 34*). Some studies suggested that pre-existent induction of CYP2E1 could play a significant role by increasing the generation of NAPQI (*20*). Hence, these thiomethyl metabolites have the potential to be used as diagnostic signals to study susceptibility to hepatotoxicity related to inter-individual production of NAPQI in these types of clinical investigations.

The current study presents limitations inherent to the production of HRMS-based semi-quantitative data. Hence, these data cannot be used to extrapolate the accurate percentage of excretion of the thiomethyl metabolites in APAP metabolism. Nevertheless, our quality control data provide confidence for the sample-to-sample comparison made to study the kinetics of the appearance of individual metabolites at different time points. The comparison of the relative contributions of APAP metabolites based on their analytical responses is therefore relevant for application as clinical markers given that similar platforms (LC-MS) and ion sources (i.e. electrospray ion sources) are usually used for APAP biomonitoring.

We also demonstrate the utility of HRMS discovery-based approach (sometimes referred to as pharmacometabolomics in clinical applications (*35, 36*)) to advance a step forward towards precision and personalized medicine and uncover new metabolites markers that can be used to provide better typing of patients related to the kinetic of elimination of a drug or to identify inter-individual susceptibility to develop drug-induced liver injury, as it is the case here with APAP. Future steps include the synthesis of pure standards for S-CH_3_-APAP-S and SO-CH_3_-APAP-S that are not currently commercially available and high-throughput targeted methods. Further validations in terms of specificity, sensitivity, and reproducibility in larger populations (including children as well) will be required for these new markers before applications in routine clinical practice.

Besides clinical applications, further toxicological investigations would also help to understand if the thiomethyl shunt acts only as a pathway for NAPQI detoxification and excretion, or if it can contribute to liver or renal toxicity in case of APAP overdose through the generation of intermediates with highly reactive sulfur-containing fragment as observed with environmental contaminants (*e.g.* trichloroethylene) (*13*). In one particular PW (*i.e.*, PW 19), unusually high levels of thiomethyl metabolites, and their intermediates were detected. Non-conjugated S-CH_3_-APAP and potential reactive intermediates (SH-APAP detected as glucuronide conjugate) were detected in the urine of this hyper-thiomethyl producer. The toxicity related to the generation of these reactive thio-intermediates remains unclear, in particular in the case of acute intoxication, and has never been studied so far in the case of APAP. Moreover, S-CH_3_-APAP was previously described as being equipotent with aspirin and more potent than APAP as an analgesic in the mouse writhing test, even though its duration of action was much shorter than the parent compound even at twice the dose (*37*). Hence, it is not clear how this S-CH_3_-APAP metabolite could contribute to the analgesic effects when produced in large quantities in humans.

In conclusion, this study provides new insights into the detoxification pathway and the elimination of the reactive NAPQI after APAP intake at therapeutic doses in humans. In particular, it highlights the role of the thiomethyl shunt pathway to form thiomethyl metabolites that could be used in the future for a better typing of patients related to the kinetic elimination of APAP and to assess inter-individual susceptibility to APAP-induced hepatotoxicity.

## MATERIALS AND METHODS

### Study design

Urine samples from one interventional study and three observational studies were used. The interventional study was designed as a longitudinal exposure to APAP over four days with an administration of 1 g of APAP on the third day. Urine samples were collected +1 h, +2h (urine only), +4h, +6h after intake, and in the morning of the subsequent two days (+24 and 48 h, urine and blood plasma). The study recruited 4 healthy men aged 30–60 years at the Department of Pharmacy and Department of Biology, University of Copenhagen. The study protocol was in compliance with the Helsinki Declaration and was approved by the Regional Scientific Ethical Committees of Copenhagen in Denmark (Protocol nr.: 17003845; and as part of trial H-17002476). All individuals provided oral informed consent to participate in the study.

The longitudinal observational study involved five pregnant women. Urine samples of these women were collected at the hospital at different time points per day, after one (pregnant woman 1), two (pregnant women 2, 3 and 4), or four (pregnant woman 5) APAP intakes. No intervention was performed for ethical reasons, and differences in dosage (500 mg vs 1g), brands (Doliprane, Dafalgan, Paracetamol Mylan), and co-administration with other pharmaceuticals were observed between intakes and pregnant women. Overall, 57 urine samples were collected, corresponding to 10, 9, 13, 13, and 12 urine collections per day for pregnant women from 1 to 5, respectively. This study was approved by the appropriate ethical committees [CPP (Comité de Protection des Personnes Sud-Est) and CNIL (Commission Nationale de l’Informatique et des Libertés).

The cross-sectional observational study included 20 pregnant women from the PELAGIE birth cohort (*38*). In this study, 10 pregnant women who declared no APAP intake versus 10 pregnant women who declared pharmaceutical consumption (including APAP) over the last 3 months were selected. These women were asked to collect, at home, the first-morning void urine sample into 10-ml vials (containing nitric acid to prevent bacterial multiplication) and to complete a questionnaire including drug intake. They returned the questionnaire and the urine sample to the research laboratory by local mail in a self-addressed stamped package. This study was approved by the appropriate ethical committees indicated above CPP and CNIL (Biological Research & Collection BRC Identification number: 2020-A00478-31).

The third observational study involved 13 individuals (11 women and 2 men) who were admitted to hospital for suspicion of acute APAP intoxication. The protocol of antidote treatment consisted of NAC administration at 150, 50 and 100 mg/kg body weight at +1h, +2-5h and +6-22h after hospitalization, respectively. Information of age of patients, Alanine transaminase (ALT) and Aspartate aminotransferase (AST) measurements, percentages of total protein and factor V is provided in table S6. Informed consent was obtained from all participants in the study and the study was approved by the Scientific Ethical Committee of Rennes Hospital in France (Avis n° 22.185).

### Targeted analyses of APAP in urine samples

Targeted analyses were used to analyze urine samples from the cross-sectional observational study to identify PW with APAP concentrations within therapeutic doses. More information on the sample preparation is provided in the Supplementary information. Briefly, extracts were analyzed for APAP after glucuronide deconjugation using a LC-MS/MS. A 1.7 µm C18 column, and the mobile phases were water and acetonitrile with 0.1% formic acid. The samples were analyzed on a Shimadzu UFLC system (Shimadzu Corporation, Kyoto, Japan) coupled to a QTRAP5500 (triple quadrupole linear ion trap mass spectrometer) equipped with a TurboIonSpray source (AB Sciex, Framingham, MA, USA). The samples were analyzed in duplicates. Excellent linearity was seen for the calibration standards ranging from 0 to 1000 ng/mL in acetonitrile/water (50:50). The correlation coefficient (r^2^) observed was above 0.994. The LOD was 2 ng/mL.

### Non-targeted analyses

#### UHPLC-ESI-QTOFMS analyses

Urine samples and medium samples from the *in vitro* liver models and the enteric bacteria were analyzed using HRMS-based non-targeted methods as described in David et al. (2021) (*11*). Briefly, urine samples (500 µl), HepaRG cell and hepatocyte samples (300 µl), and microbiota samples (150 µl) were diluted to 1 ml of HPLC grade water, acidified with 1% formic acid and extracted using Strata-X SPE. All sample extracts were profiled using an Exon UHPLC system (AB Sciex, USA) coupled to an AB Sciex X500R Q-TOF-MS system (Sciex technologies, Canada), equipped with a DuoSpray ion source. As a first step, all the samples were analyzed in a full-scan experiment (50–1100 Da) in both – and + ESI modes. MS/MS mass fragmentation information for chemical elucidation was obtained by further analysis of selected samples in sequential window acquisition of theoretical mass spectrum (SWATH) data-independent acquisitions. Quality controls for HRMS-based analyses are described in the Supplementary information.

### Data processing and annotation

SciexOS and vendor software MarkerView 1.3.1. (AB Sciex, USA) were used for data acquisition control and initial data preprocessing for quality control. The obtained LC-MS raw files (wiff2 format) were converted into mzXML format using MSConvert GUI (Palo Alto, CA, USA) using the Proteowizard open-source software (*44*). Converted data were then imported to MATLAB computer and visualization environment (Release 2021b, The Mathworks, Inc., Natick, MA, USA) and analyzed with MSroi (*45*) chemometrics strategy using the MATLAB MSroi app (*45*). This approach was employed for data compression, feature detection, and filtering (more information is provided in the Supplementary information). Multivariate and univariate analyses were performed in a MATLAB environment in order to identify APAP metabolites after APAP, APAP-Cys, NAC-APAP, and APAP-SG exposures as described in the supplementary information. The identities of the expected APAP metabolites were determined from accurate mass, isotopic fit, and fragmentation data obtained from SWATH acquisition and from comparison with standard compounds when available or spectra available in online libraries or the literature (*11*). In all cases, metabolite identification was based on recommendations by Schymanski et al. (table S9).

### Accurate integration of identified and annotated markers

Peak integration of all APAP metabolites was performed manually using the Sciex OS Analytics tool. The integrated peak area of individual markers was normalised by dividing the area of each metabolite by the total peak area of the chromatogram, measured with MarkerView. MATLAB (Release 2021b, The Mathworks, Inc., Natick, MA, USA) was used to generate plots.

### Cell culture and treatment of HepaRG cells and human hepatocytes

Highly differentiated human HepaRG cells from passages 13 to 16 were cultured as described (*39*). Briefly, cells plated at a density of 2 × 10^5^ cells/cm^2^ in 12-well tissue culture-treated polystyrene plates, were first grown for 2 weeks in Williams’ E medium (Invitrogen, Cergy-Pontoise, France) supplemented with 10% (vol/vol) fetal calf serum (FCS) (Cytiva, Velizy-Villacoublay, France), 100 IU/ml penicillin, 100 μg/ml streptomycin, 5 μg/ml bovine insulin (Sigma–Aldrich, Saint-Quentin-Fallavier, France), 2 mM glutamine (Invitrogen) and 50 µM hydrocortisone hemisuccinate (Upjohn, Paris La Défense, France). Cells were next cultured for an additional 2 weeks-period in the same medium supplemented with 2% (vol/vol) dimethyl sulphoxide (DMSO), in order to get a full differentiation status of the cells (*40*) before their use for metabolism activity assay. This differentiated status of DMSO-treated HepaRG cell cultures was routinely checked by their phase-contrast microscopy analysis, demonstrating the presence of hepatocyte-like islands and the formation of bile canaliculi, as previously reported (*39*). For establishing human hepatocyte cultures, fresh or cryopreserved human hepatocytes, prepared from human liver fragments, were obtained from Biopredic International (Saint-Grégoire, France). Cells were directly (for fresh hepatocytes) or after thawing (for cryopreserved hepatocytes) suspended at a concentration of 1 x 10^6^ cells/ml in a seeding medium composed of Williams’ E medium, supplemented with 10% (vol/vol) FCS, 5 µg/ml bovine insulin, 100 IU/ml penicillin, 100 µg/ml streptomycin, and 2 mM glutamine. Cells were then seeded on 24-well tissue culture-treated polystyrene plates at a density of 2 x 10^5^ cells/cm^2^. After 24 h, the medium was discarded, and hepatocytes were cultured in a maintenance medium composed of seeding medium, supplemented by 50 µM hydrocortisone hemisuccinate and 2% (vol/vol) DMSO, as previously described (*41*). The culture medium was routinely renewed every 2-3 days and the cells were used for metabolism activity assays after one week of culture.

For cell treatment by APAP precursors, HepaRG cells and primary human hepatocytes were first incubated with 50 µg/ml of APAP or APAP-cys in the maintenance medium described above, during 0.5 h, 2 h, 6 h, 24 h, or 48 h, to optimize the incubation time required to detect APAP metabolites. Cells were next exposed for 24 h to 50 µg/ml of APAP-Cys, APAP-SG, or NAC-APAP.

### Enteric bacteria

Five gut bacterial strains were selected including *Escherichia coli* (Ecoli), *Enterococcus faecalis* (efae), *Agathobacter rectalis* (arec), *Phocaeicola vulgatus* (pvul) and *Bifidobacterium longum* (blon). The bacterial strains used in this study were selected to be representative of each phylum present in the gut microbiome (*42, 43*). *Agathobacter rectalis* DSM 17629, *Phocaeicola vulgatus* DSM 1447, *Bifidobacterium longum subsp*. Longum DSM 20219 were provided by DSMZ Gmbh, Braunschweig, Germany. *Escherichia coli* DSM 1576 and *Enterococcus faecalis* ATCC19433 were provided by CIP (Institut Pasteur Collection, Paris, France). Cultures were performed in GAM medium (Gifu anaerobic broth, Fisher scientific, Illkirch, France), at 37°C with agitation under anaerobic conditions generated by Genbag anaer (Biomerieux, Marcy l’Etoile, France). Growth kinetics were performed for each strain to identify the time required to reach the exponential phase. For this purpose, a multiskan FC spectrophotometer (Thermo Scientific, Illkirch, France) was used to measure the optical density at 620 nm.

For exposure experiments, culture growth was stopped during the exponential phase in order to prepare lysates or strains. For lysates preparation (*31*), cultures were centrifuged (10,000 × g, 15 minutes), washed twice with potassium phosphate buffer (100 mM, pH 7.0), resuspended in borate buffer (50 mM, pH=8.4), and broken by mechanical lysis with glass beads in a tissue lyser (Qiagen, Courtaboeuf, France) 10 minutes, 50 Hz. The extracts were obtained after removing the beads by centrifugation. The lysates were exposed to 50 µg/ml of APAP and APAP-Cys for 20 minutes. For strain preparation, cultures were centrifuged at 8000g for 3 min, washed in PBS (pH=7), and resuspended in the same buffer. The E coli strain was exposed to 50 µg/ml of APAP and APAP-Cys for 30 minutes, 2h, 6h, 24h. Then all strains were exposed to 50 µg/ml APAP and APAP-Cys for 24h.

### Statistical Analysis

Univariate analyses were performed to compare fold changes, calculate the areas under the curve (AUC) and determine statistical significance. AUCs were calculated using trapezoidal numerical integration (trapz function) (*46*) from MATLAB. Statistics Toolbox^TM^ from MATLAB (Eigenvector Research Inc., Wenatchee, WA, USA) was used to assess statistical significance using a one-way ANOVA t-test, considering *P* < 0.001 statistically significant, followed by a multiple comparison test (multcompare) to perform pairwise comparisons, based on the Tukey-Kramer critical value (*47*).

### List of Supplementary Materials

#### Materials and Methods

Fig. S1 Kinetics of formation (t= 0, 1, 2, 4, 6, 24 and 48 h) of APAP metabolites in urine samples of the controlled study performed in 4 men.

Fig. S2 Kinetics of formation (t= 0, 0.5, 2, 6, 24 and 48h) of APAP metabolites in HRG cells and Hepatocytes after exposure to APAP and APAP-Cys, including a zoom of metabolites showing low intensity signals.

Fig. S3 APAP metabolites produced in 5 microbiota lysate strains after exposure to APAP and APAP-Cys.

Table S1. AUCs of APAP metabolites for the five pregnant women of the longitudinal observational study and normalised UHPLC areas.

Table S2. AUCs of APAP metabolites in the controlled study of 4-men and normalised UHPLC areas.

Table S3. Drug administration for the five pregnant women of the longitudinal observational study.

Table S4. Normalised UHPLC areas and “Total APAP” measured by ELISA in the urine samples of the 20 pregnant women of the cross-sectional observational study (cohort PELAGIE).

Drug administration for the five pregnant women of the longitudinal observational study.

Table S5. Names, formulas, m/z values, retention times and areas of the 7 APAP standards used in the study.

Table S6. Information on APAP, ALT/AST concentrations in blood samples of 13 individuals suspected of APAP acute intoxication

Table S7. Normalised UHPLC areas, “Total APAP” measured by ELISA and correlations among them in blood samples of the 13 intoxicated patients.

Table S8. List of standards used for the identification of APAP metabolites and the 15 labeled internal standards (IS) spiked in samples for the non-targeted analyses and their respective suppliers.

Table S9. Annotation parameters for APAP metabolites according to Schymanski et al. (2014) Data file S1 or Data files S1 to Sx (Excel files)

References (*11*) is cited in Supp Material

## Acknowledgments

We thanks Thibaut Léger and Christine Monfort for technical assistance.

## Author contributions

Each author’s contribution(s) to the paper should be listed [we encourage you to follow the CRediT model]. Each CRediT role should have its own line, and there should not be any punctuation in the initials.

Conceptualization: AD, EG

Methodology: AD, EG, AG, HS, JC, MLV, PLC, CC, TG, OF, AMK, CL, BJ

Investigation: AD, EG, AG, MLV, HS, TG, OF, DK Visualization: AD, EG, AG, HS

Funding acquisition: AD, RB, GWM Project administration: AD, RB, GWM Supervision: AD

Writing – original draft: AD, EG

Writing – review & editing: AD, EG, AG, HS, JC, MLV, PLC, CC, TG, OF, AMK, CL, DK, RB, GWM

