## Supplementary Figures for "Paracetamol/acetaminophen hepatotoxicity: new markers for monitoring the elimination of the reactive N-Acetyl-p-benzoquinone imine"

**Targeted analyses of APAP in urine samples**

Targeted analyses were used to analyse urine samples from the cross-sectional observational study to identify PW with APAP concentrations within therapeutic doses. Twenty µL of urine were transferred to a glass insert (Teknolab Sorbent, Kungsbacka, Sweden) of a 96-well Rittner plate (Teknolab Sorbent). Ten µL of ammonium acetate (pH 6.5) and 0.01 mL glucuronidase (*Escherichia coli*) were added, and the solution was incubated at 37 °C for 90 min. After addition of 0.025 mL of MilliQ water and 0.3 mL of internal standard, the plates were centrifuged for 10 min at 3000 rpm, prior to injection. Extracts were analysed for APAP after glucuronide deconjugation using LC-MS/MS. A 4 µm C18 column (2.1 mm i.d. × 50 mm, Genesis Lightening) was used before the injector to reduce the interference of contaminants during the mobile phase. A 1.7 µm C18 column (2.1 mm i.d. × 100 mm; Fortis Technologies) was used for separation, and the mobile phases were water and acetonitrile with 0.1% formic acid. The samples were analysed on a Shimadzu UFLC system (Shimadzu Corporation, Kyoto, Japan) coupled to a QTRAP5500 (triple quadrupole linear ion trap mass spectrometer) equipped with a TurboIon Spray source (AB Sciex, Framingham, MA, USA), in duplicates. All runs included at least 10 blank samples, used for a chemical noise subtraction step for all biological samples. Excellent linearity was seen for the calibration standards ranging from 0 to 1000 ng/mL in acetonitrile/water (50:50). The correlation coefficient ($r^2$) observed was above 0.994. LOD and LOQ were determined by analysis of 10 different blank toluene samples spiked with internal standards. The LOD was calculated as three times the standard deviation of the ratio between the peak area at the analyte retention time and the peak area of internal standard, divided by the slope of the calibration line. The determined LOD was 2 ng/ml.

**Non-targeted analyses**

**Chemicals**

The list of standards used for the identification of APAP metabolites and the 15 labelled internal standards (IS) spiked in samples for the untargeted analyses and their respective suppliers are provided in Supplementary Table 7. All solvents were high-performance liquid chromatography grade, purchased from Biosolve Chime (Dieuze, France). Strata-X Polymeric Reversed Phase cartridges (200 mg, 3 mL) were supplied by Phenomenex (Le Pecq, France).

**Quality control**

In order to evaluate and control potential background contaminants, one workup sample (i.e., extraction with HPLC grade water instead of sample) per analytical batch was prepared and injected. Quality control (QC) samples, consisting on pools of all biological samples, were injected all along the analytical batch in order to monitor for UHPLC-ESI-TOF-MS repeatability and sensitivity and to normalize concentrations among metabolites among sample sets. To ensure no presence of remaining traces in the UHPLC system from previous batches that could affect subsequent analytical runs, solvent blank samples (acetonitrile/$H_2O$ (20:80)) were also injected within the analytical batch. Sample batch order of injection was as follows: first injections corresponded to blank samples (workup and solvent), followed by the injection of a QC sample six times consecutively, and then all samples from the batch, randomly distributed in sets of 5, interspersed each time with a periodic QC to monitor the analytical sensitivity and repeatability.

**Data processing and annotation.**

Converted data were then imported to MATLAB computer and visualization environment (Release 2021b, The Mathworks, Inc., Natick, MA, USA) and analysed with MSroi *(45)* chemometrics strategy using the MATLAB MSroi app *(45)*. This approach was employed for data compression, feature detection and filtering. Shortly, MSroi enables significant data compression in the spectral dimension without losing their instrumental spectral accuracy. The main parameters used for the ROI procedure in this work were a mass error tolerance of 10 ppm, a minimum threshold value of 1000, and a minimum occurrence value of 10 seconds, and ROIs m/z values were calculated with the median of the m/z values determined for each chromatographic peak. The outcome of the ROI approach consists of a vector including the list of relevant *m/z* ROI values (according to the previously mentioned parameters), and a data matrix with the MS intensities at the selected ROIs (for all considered retention times and samples). The graphical interphase allows the visualization of the ROI features. The ROI vectors and MSroi matrix containing information of the features were used for multivariate and univariate analysis in MATLAB environment in order to identify APAP metabolites after APAP, APAP-Cys, NAC-APAP, and APAP-SG exposures. Principal component analysis (PCA) were performed using the PLS Toolbox (Eigenvector Research Inc., Wenatchee, WA, USA) as a first step to examine QC clustering, sample separation and to identify any analytical or biological outliers. Partial least squares-discriminant analysis (PLS-DA) was then performed and by the use of VIP scores, the compounds showing higher VIP score were selected as potential biomarkers of exposure.

The discriminating markers between exposed and control groups identified through multivariate analyses were included in a suspect list. The suspect list included all the APAP metabolites found in a previous study of the same authors *(11)* and also included other metabolites available in the literature *(27)*. As explained in a previous study, the identities of the expected APAP metabolites were determined from accurate mass, isotopic fit, fragmentation data obtained from SWATH acquisition and from comparison with standard compounds when available or spectra available in online libraries or the literature *(11)*. In all cases, metabolite identification was based on recommendations by Schymanski et al. (Supplementary Table S9).

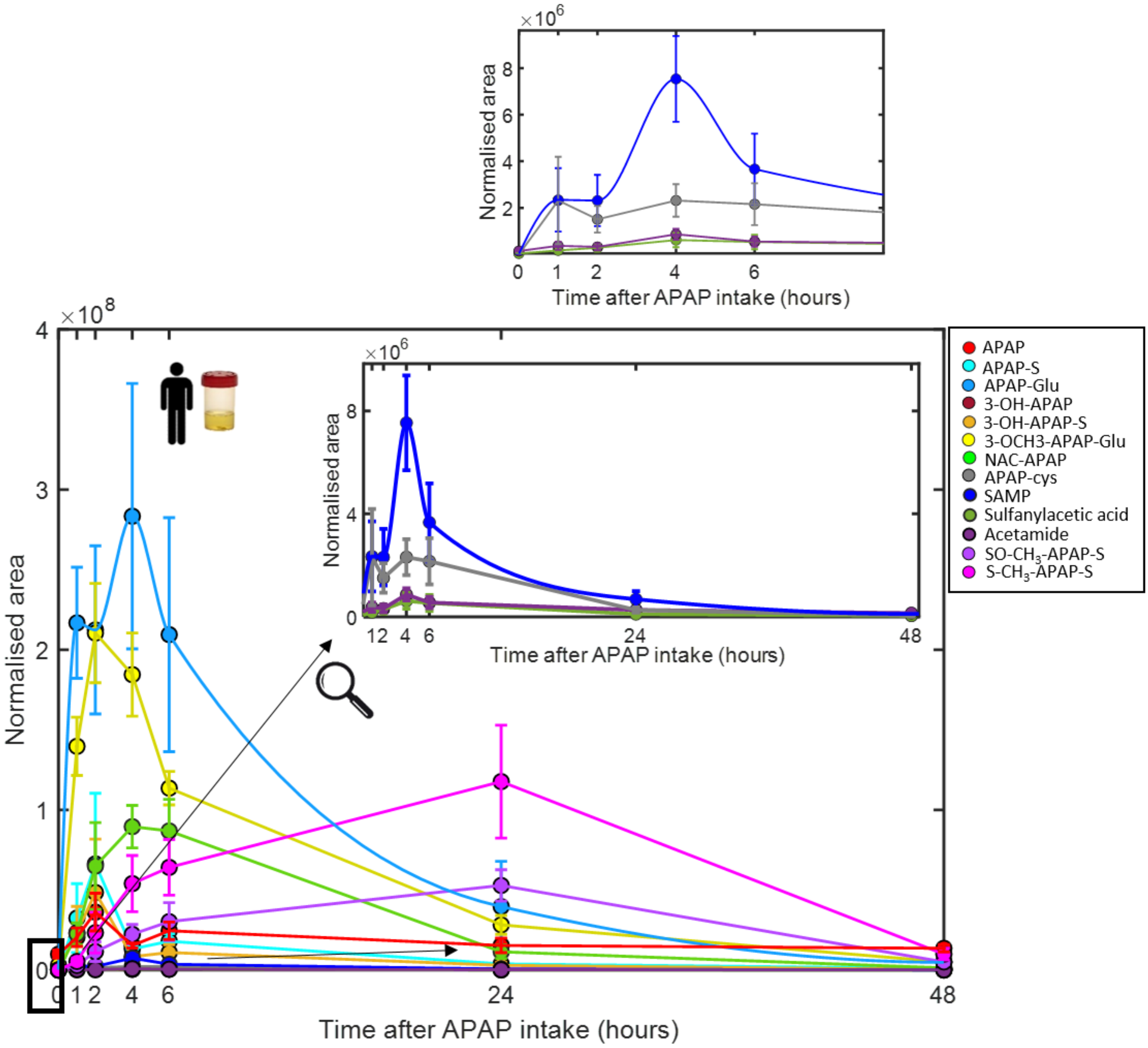


**Figure S1**. Kinetics of formation of APAP metabolites in the urine of 4-men (mean ± SEM) before APAP intake (baseline, n=12) and at different time points (0, 1, 2, 4, 6, 24 and 48h) after 1g APAP intake (control study, *(11)*).

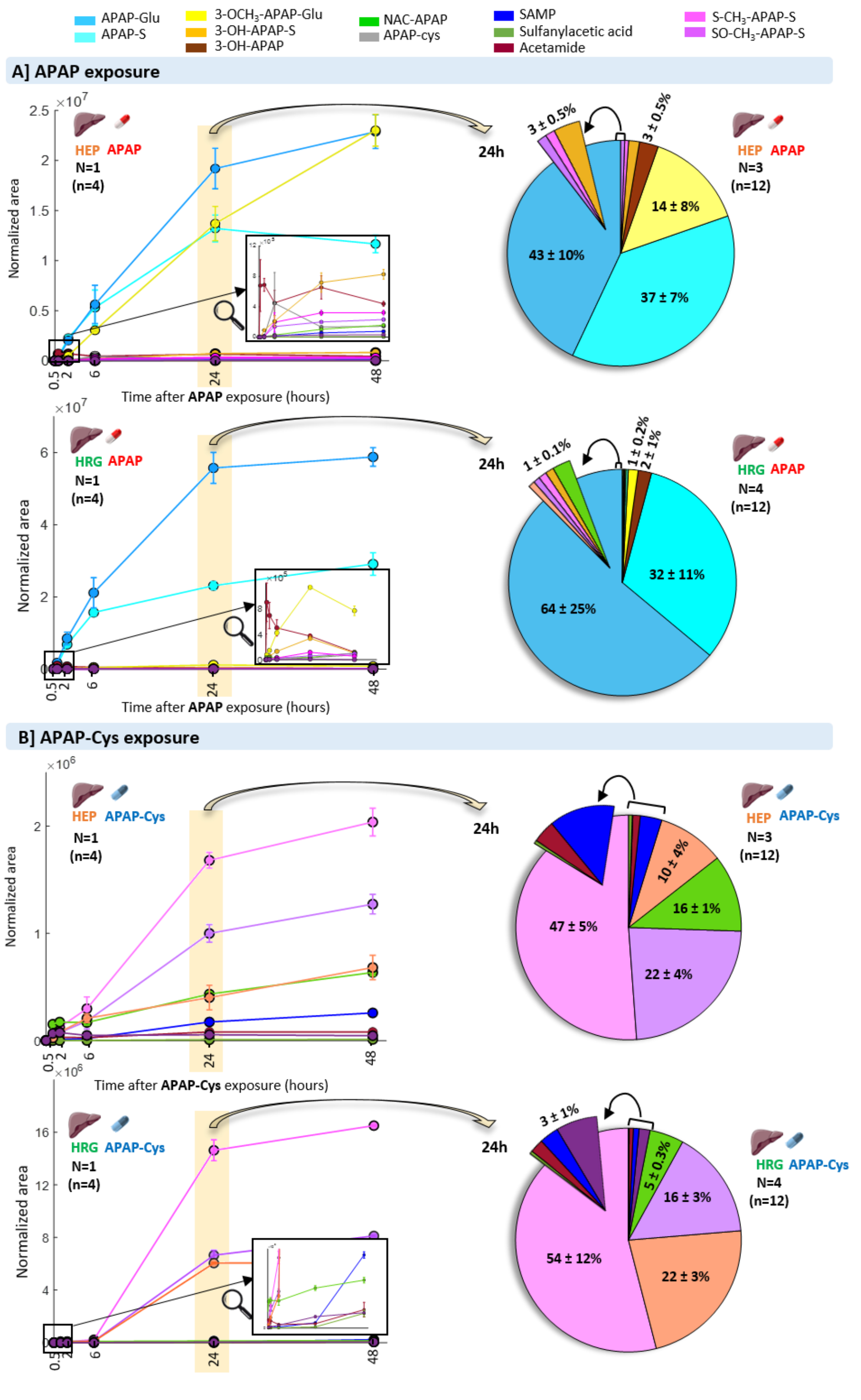


**Figure S3**. Figure 3 in the main text including the zoom of the low signal metabolites.

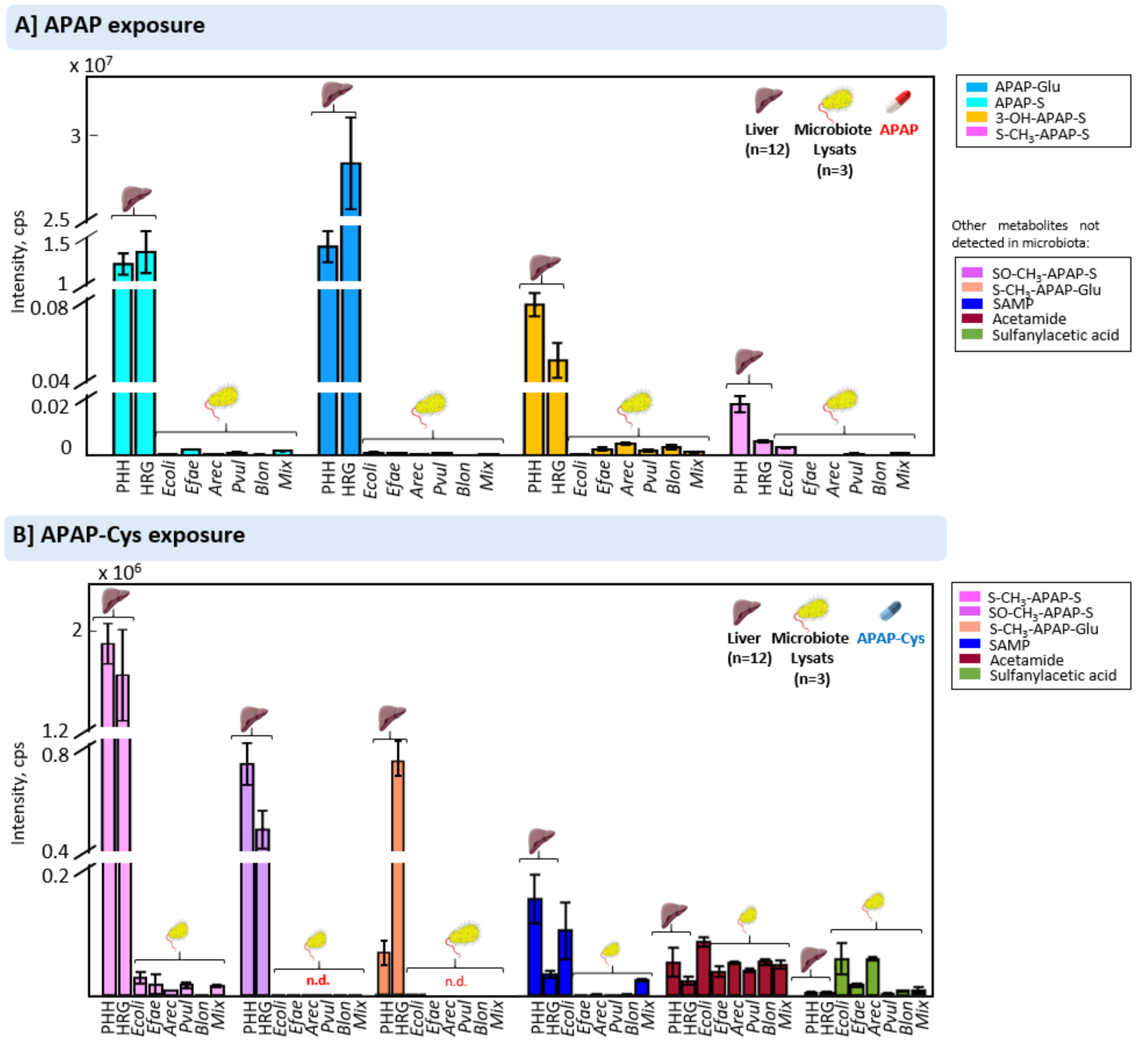


**Figure S3** Acetaminophen metabolites produced HepaRG cells, Hepatocytes and 5 types of intestinal microbiote lysats after 20 min exposure to A) free acetaminophen (APAP) and to B) 3-(Cystein-S-yl)acetaminophen (APAP-Cys) both at a concentration of [50 µg/ml]. Abbreviations: *Escherichia coli* (Ecoli), *Enterococcus faecalis* (Efae), *Agathobacter rectalis* (Arec), *Phocaeiola vulgatus* (Pvul), *Bifidobacterium longum* (Blon) and mixture of the five strains (Mix). n=total number of replicates.
